# GABAergic cell loss in mice lacking autism-associated gene *Sema6A*

**DOI:** 10.1101/663419

**Authors:** Karlie Menzel, Gábor Szabó, Yuchio Yanagawa, Turhan Cocksaygan, Céline Plachez

## Abstract

**Background:** During brain development, a multitude of neuronal networks form as neurons find their correct position within the brain and send out axons to synapse onto specific targets. Altered neuronal connectivity within these complex networks has been reported in Autism Spectrum Disorder (ASD), leading to alterations in brain function and multisensory integration. Semaphorins (also referred to as Semas), a large protein family of about 30 members, have been shown to play an important role in neuronal circuit formation and have been implicated in the etiology of ASD. The purpose of the current study is to investigate how *Sema6A* mutation affects neuronal connectivity in ASD. Since *Sema6A* is involved in cell migration, we hypothesized that during brain development the migration of GABAergic interneurons is affected by the loss of *Sema6A* gene, leading to alterations in Excitatory/Inhibitory (E/I) balance.

**Methods:** *Sema6A* transgenic mice were crossed with either GAD65-GFP mice or GAD67-GFP mice to allow for both a reliable and robust staining of the GABAergic interneuron population within the *Sema6A* mouse line. Using histological techniques we studies the expression of interneurons subtypes in the Sema6A mutant mice.

**Results:** Analysis of *Sema6A* mutant mice crossed with either GAD65-GFP or GAD67-GFP knock-in mice revealed a reduced number of GABAergic interneurons in the primary somatosensory cortex, hippocampus, and reticular thalamic nucleus (RTN) in adult *Sema6A* mutant mice. This reduction in cell number appeared to be targeted to the Parvalbumin (PV) interneuron cell population since neither the Calretinin nor the Calbindin expressing interneurons were affected by the *Sema6A* mutation.

**Limitations:** Although the use of animal models has been crucial for understanding the biological basis of autism, the complexity of the human brain can never truly be replicated by these models.

**Conclusions:** Taken together, these findings suggest that *Sema6A* gene loss affects only the fast spiking-PV population and reveal the importance of an axon guidance molecule in the formation of GABAergic neuronal networks and provide insight into the molecular pathways that may lead to altered neuronal connectivity and E/I imbalance in ASD.

## BACKGROUND

Leo Kanner was the first to describe early infantile autism in 1944 [1] and since then tremendous effort has been made to not only understand the causes of Autism Spectrum Disorder (ASD) but also to improve early diagnosis. While it appears that the causation of autism is very complex, one can learn from studying abnormalities in brain structure or function. Magnetic Resonance Imaging (MRI) as well as histological techniques using brain tissue from individuals with autism have provided invaluable data in the quest to elucidate brain differences occurring in ASD. Indeed many studies that have examined brain anatomy at a structural and/or cellular level have reported an increased volume for total brain, parieto-temporal lobe, and cerebellar hemispheres in ASD [2]. Brain growth and connectivity differences are apparent early in brain development [3, 4] and a recent review by Donovan and Basson [5] provided evidence that brain growth is altered in ASD. Neuropathological studies revealed altered neurogenesis and defective neuronal migration resulting in focal dysplasia in the cortical, hippocampal, and cerebellar brain areas [6, 7]. All these changes in neuronal organization suggest that ASD brain differences take place early in brain development. In addition, recent human and animal studies have suggested that many psychiatric disorders, including ASD, are characterized by an imbalance between excitation and inhibition (E/I) in the brain affecting normal brain activity [8] [9] [10] [11] [12]. Excitatory glutamate pyramidal neurons and inhibitory GABAergic interneurons are the main components of cortical neural circuits and GABAergic interneurons regulate excessive excitation of pyramidal neurons [11]. Differences in brain growth and connectivity occurring early in brain development, coupled with E/I imbalance, suggest that either the production, migration, cell-type specification, or maturation of GABAergic interneurons may be altered in ASD. Both excitatory and inhibitory neurons are guided by a combination of chemo-attractive and repulsive cues [13] [14] [15, 16]. Hussman et al. 2011 [17], in a genome-wide association study, reported autism candidate genes with known roles in neurite outgrowth and guidance including the *Semaphorin-6A* (also called *Sema6A)* gene. Semaphorins (also referred to as Semas) constitute a large protein family of 30 members [16, 18] and their role in the nervous system has been widely studied. Acting as guidance molecules, Semas have been shown to be implicated in many processes of development. More importantly, some members of the Semaphorin family have been shown to play an important role in neuronal circuit formation and have been implicated in the etiology of ASD [19–24]. Recent studies have shed light on the importance of *Sema6A* as a candidate gene in ASD [17, 24–27]. Increasing evidence suggests a role for *Sema6A* gene in ASD, as it was implicated as an autism candidate gene in both genome-wide prediction [17, 26] (prediction rank can been seen at http://asd.princeton.edu) and ASD IQ gene analysis [25]. Data obtained from two studies of *Sema6A* mutant mice also suggest a role for *Sema6A* in ASD [27, 28]. The first study revealed the role of *Sema6A* in the formation of both limbic and cortical connectivity as *Sema6A* mice displayed cellular disorganization and dysconnectivity [24]. The second study explored fundamental levels of behavior and reported a dysfunction in both motivational and motoric processes [27]. Taken together these studies revealed the potential for using *Sema6A* mutant mice as an animal model to study dysfunction in brain neuronal networks, as they displayed brain alterations similar to those seen in ASD, and the loss of a guidance molecule could also lead to disrupted E/I balance. Therefore the present study focuses on the contribution of *Sema6A* toward GABAergic interneuron migration during brain development as an underlying cause of ASD. Taking advantage of genetically engineered mouse lines, we observed a reduction in the number of GABAergic cells in *Sema6A* mutant mice, and found that the loss of *Sema6A* affected more specifically the calcium-binding parvalbumin interneuron population. These results reveal a role for *Sema6A* in the formation of neuronal circuits in cortical, hippocampal, and reticular thalamic nucleus areas: key brain areas where alterations have been reported in individuals with ASD.

## METHODS

### Animals

All experimental procedures were performed in compliance with protocols approved by the Institutional Animal Care and Use Committees (IACUC) from the Hussman Institute for Autism and University of Maryland, Baltimore (UMB). Animals were housed at UMB, an AAALAC accredited facility. Animals were subjected to a standard 12 hr light/dark cycle and *ad libitum* access to water and food. *Sema6A* mutant mice were obtained from Dr. Giovanna Tosato, laboratory of cellular oncology at NCI/CCR (NIH), with the permission of Dr. Kevin Mitchell, Trinity College Dubin. Generation of the *Sema6A* mouse line was previously described by Leighton et al., 2001 [29] and Kerjan et al., 2005 [30]. GAD65-GFP transgenic mouse strain was used for this project. In this mouse line, green fluorescent protein (GFP) is expressed under the control of the GAD65 promoter. This mouse line was first developed by Gábor Szabó [31] in the Medical Gene Technology Unit, Institute of Experimental Medicine in Budapest, Hungary. GAD67-GFP transgenic mouse strain was also used in the project, in which GFP is expressed under the control of the endogenous GAD67 promoter [32]. This mouse line was developed by Yuchio Yanagawa [33] in the Department of Genetic and Behavioral Neuroscience, Gunma University Graduate School of Medicine, Japan. For immunohischemistry purposes, animals were anesthetized with isoflurane and then subjected to transcardiac perfusion with 0.9% of sodium chloride (NaCl) followed by 4% paraformaldehyde (PFA). Tissue was stored in 4% PFA at 4°C until dissection.

### PCR

DNA from 2mm tail clippings were extracted with QuickExtract kit (Epicentre). *Sema6A* genotype was determined using the following primers: 5’-gagatgcacagctaacttctggtg-3’ (wild-type allele forward primer), 5’-ttgaagcctgctcttagtggctcc-3’ (reverse primer), and 5’-gctaccggctaaaacttgagacct-3’ (mutant allele reverse primer). This amplified a 990-bp product for the mutant allele and a 1.43-bp product for the wild-type allele. PCR was completed using RedExtract ReadyAmp Taq (Sigma, SIG-R4775) with the following cycle conditions (29 cycles): 95°C for 30 seconds, 64°C for 30 seconds, and 72°C for 2 minutes. PCR were run on either a T100 (Bio-rad) or a SimpliAmp (Life Technologies) thermal cycler and were imaged using ChemiDoc Touch Imaging System (Bio-rad).

### Immunohistochemistry

Brains were dissected, cryo-protected with 30% sucrose in 1X PBS (phosphate buffer saline), embedded in Optimal Cutting Temperature (OCT – VWR) and sliced at 40µM (Leica, cryostat). Sections were washed in 1X PBS and then blocked for 30 minutes at room temperature in blocking solution (BS) [BS contains 1X Tris Buffer solution (TBS)-0.3% Triton, 1% Donkey Serum, and 1% Bovine serum albumin (BSA)] followed by incubation with primary antibodies at room temperature overnight (see antibody table for details on the primary antibodies that were used in this study). The next day, sections were rinsed in 1X TBS-0.3% Triton and incubated with the corresponding secondary antibodies for two hours at room temperature (see antibody table for details on the secondary antibodies used). For DAB (diaminobenzidine) staining, VECTASTAIN Elite ABC kit was used (Vector Laboratories, PK-6100) at room temperature for 1 hour, sections were rinsed and then incubated at room temperature in DAB (Sigma, cat# D4418).

### Antibodies

| Primary Antibodies | Source | Cat# | Concentration |
| --- | --- | --- | --- |
| Rabbit anti-Calbindin | Swant* | CB-38a | 1/100K |
| Rabbit anti-Calretinin | Swant* | 7699/4 | 1/30K |
| Rabbit anti-Calretinin | Millipore/Sigma<sup>◇</sup> | AB5054 | 1/30K |
| Guinea pig anti-Parvalbumin | Swant* | GP72 | 1/5K |
| Secondary Antibodies | Source | Cat# | Concentration |
| Biotin-SP-conjugated Donkey Anti-Rabbit IgG | JIR‡ | 711-065-152 | 1/2K |
| Biotin-SP-conjugated Donkey Anti-Guinea Pig IgG | JIR‡ | 706-065-148 | 1/2K |

\* Swant, https://www.swant.com; ◇ Millipore/Sigma: http://www.emdmillipore.com; ‡JIR: Jackson ImmunoResearch: https://www.jacksonimmuno.com.

#### Histological analysis

All images were obtained using an Axio Zoom V16 microscope (Zeiss) and NIH image J software was used for cortical thickness, cortical area and cell number analysis. Histological and cell count analyses were performed blind of genotype (wildtype, heterozygous, or knockout) and sex. Cortical thickness was measured in the primary somatosensory cortex (S1) from bregma levels −1.06 mm to −2.06 mm. Measurements of cortical area were achieved using a boundary map of the neocortical areas, including motor, somatosensory and auditory cortical areas at bregma levels −1.06 mm to −2.06 mm. Cell number analysis for both GABAergic interneurons, GAD65-GFP and GAD67-GFP, consisted of counting fluorescent GFP positive cells in the primary somatosensory cortex (S1), hippocampus and reticular thalamic nucleus at bregma levels −1.06 mm to −2.06 mm. Analysis of either Parvalbumin, or Calretinin or Calbindin cell number consisted of counting DAB positive cells in the primary somatosensory cortex (S1), hippocampus and reticular thalamic nucleus at bregma levels −1.06 mm to −2.06 mm. A minimum of 7 animals and a maximum of 12 animals were used for all the analysis. There were no significant differences between the cell counts from males and females of the same genotype therefore the data presented in this study combines cell counts obtained from either males or females of the same genotype. Histological data were analyzed using unpaired two-tailed t tests using GraphPad Prism 8.0.0 (GraphPad Software; San Diego, CA), with genotype (Control *vs* KO) as the main factor. Data are presented as mean ± SEM and statistical significance was set at p ≤ 0.05 – see figure legends for number of animal per analysis and p values.

## RESULTS

### Both cortical thickness and cortical area were unchanged in *Sema6A* mutant mice

To investigate whether *Sema6A* mutants can be used as an informative model to investigate the E/I imbalance seen in ASD, the overall brain anatomy of the *Sema6A* mutant mice was first analyzed. As previously reported by Runker et al. 2011 [24], our results showed that *Sema6A* mutant mice at P30 displayed cortical lamination defects in the auditory cortex (data not shown), however both primary somatosensory cortical (Fig. 1A and A’ compared to Fig. 1B and B’) and hippocampal areas (Fig. 1A and A” compared to Fig. 1B and B”), where our data processing of the GABAergic population focused, appeared to be unaffected by the loss of *Sema6A* gene. Analysis of the cortical thickness as well as the cortical area revealed no difference between *Sema6A* mutant and control mice at P30 (Fig.1C and D, mutant mice n=7, control mice n=7). In order to investigate the contribution of *Sema6A* toward GABAergic interneuron migration, *Sema6A* mice were crossed with either GAD65-GFP transgenic mice or GAD67-GFP knock-in mice.

**Figure 1:**
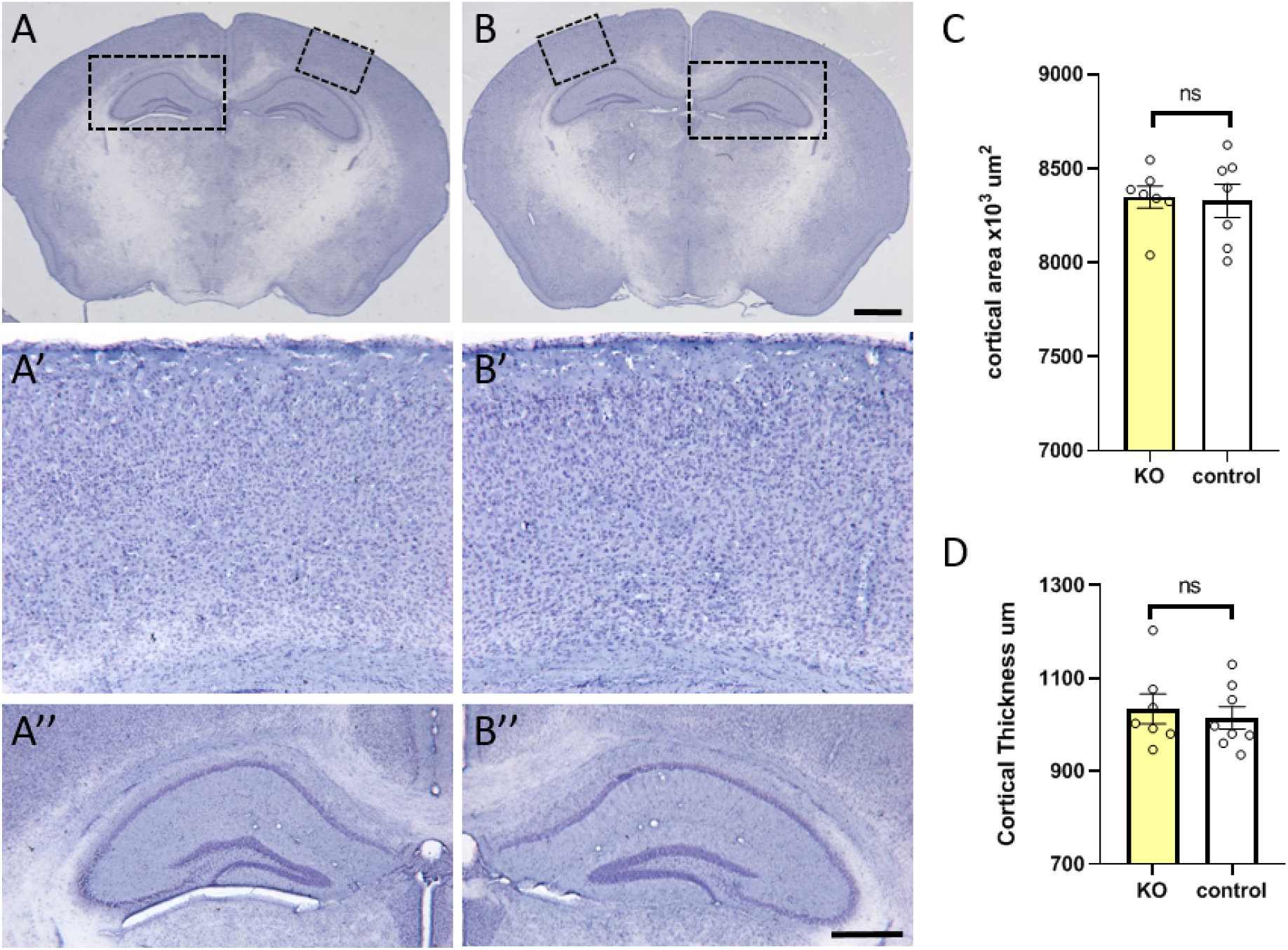
Cortical area and cortical thickness were not affected in *Sema6A* mutant mice. Hematoxylin staining was performed on P30 coronal brain slices. No anatomical differences were found between *Sema6A* mutant (A, A’, A”) and *Sema6A* control (B, B’, B”) brains. A’ and B’ are high magnification of the inserts in A and B, respectively, at cortical level. A” and B” are high magnification of the inserts in A and B, respectively, at the hippocampal level. Both the cortical area (C) and the cortical thickness (D) were measured in both *Sema6A* mutant and control mice and no significant differences were found. (ns: not significant in C and D). Scale bar in B is 1000µm for A and B; scale bar in B” is 200µm for A’ and B’ and 400µm for A” and B”.

### Reduction in GAD65-GFP positive cells in *Sema6A* KO mice

GABA synthesis from glutamate is dependent upon glutamate decarboxylase (GAD) enzyme. There are two distinct isoforms of GAD, GAD65 (65 kDa form) and GAD67 (67 kDa form), both able to synthesize GABA [34]. GAD65-GFP transgenic mouse line in which all GAD65 inhibitory neurons express GFP [35] has proven to be a very useful tool to visualize GABAergic interneurons considering the relatively low sensitivity of GABA immunoreactivity. Therefore, GAD65-GFP mice were crossed with *Sema6A* mice to allow for a reliable and robust staining of the GAD65 interneuron population within the *Sema6A* mouse line and for a better understanding of the type of cells affected by the mutation. In order to match closely the neuronal network alterations seen in adult humans with ASD, GABAergic interneuron expression was analyzed at postnatal stage 30 (P30) as it is considered an adult stage with a completely formed neuronal network. Our results showed that adult *Sema6A*/GAD65-GFP mutant mice have a reduced number of GAD65-GFP+ cells. This reduction was present in the primary somatosensory cortex (S1) and well as in hippocampal and reticular thalamic nucleus (RTN) areas of *Sema6A* mutant mice (Fig. 2 A-A’”) compared to *Sema6A* WT mice (Fig. 2 B-B’”). Cell counts revealed that GAD65-GFP+ cell number was reduced by 50% in the cortical area (Fig. 2C, mutant n=8, control n=8), 46% in the hippocampal area (Fig. 2D, mutant n=8, control n=8), and 62% in the RTN area (Fig. 2E, mutant n=8, control n=7) compared to the control means for each area. While *Sema6A* mutant mice displayed such a dramatic loss of GABAergic interneurons, there were no anatomical differences visible (Fig. 1) as measures of both cortical thickness and cortical area revealed no differences between *Sema6A* mutant and control (Fig. 1C and 1D) mice.

**Figure 2:**
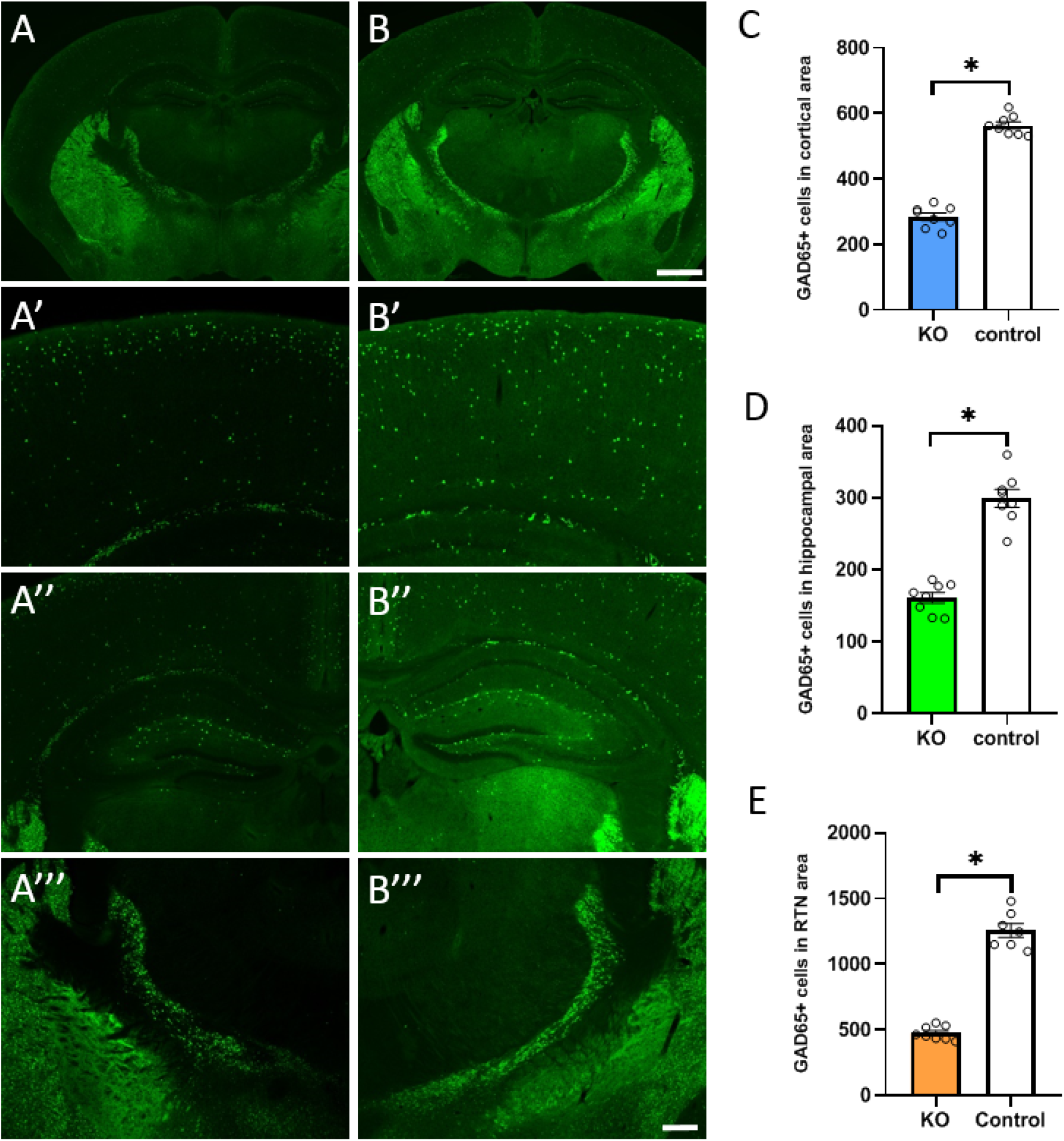
*Sema6A* mutant mice display a decrease in GAD65-GFP positive cells. *Sema6A* mutant mice were crossed with GAD65-GFP mice to allow for reliable labeling of the GAD65 interneurons population (green staining on panels A-A”’, B-B’”). Panels A’-A’” shows GAD65-GFP positive cells (in green) in a *Sema6A* mutant mouse at P30. A’, A” and A’” are higher magnification views of the primary somatosensory cortex, hippocampus and RTN, respectively. Panels B, B’, and B’” show GAD65-GFP positive cells in a *Sema6A* control mouse at P30. B’, B” and B’” are higher magnification views of the cortex, hippocampus and RTN, respectively. Panels C, D and E represent GAD65-GFP positive cell count with a reduction of GAD65 positive cells seen in *Sema6A* mutant mice at P30 in the cortical area (C, blue bar: KO, n=8, white bar: WT, n=8, p value<0.0001), hippocampal area (D, green bar: KO, n=8, white bar: WT, n=8, p value<0.0001) and reticular thalamic nucleus (E, orange bar: KO, n=8, white bar: WT, n=8, p value<0.0001). Scale bar in B is 1000µm for A and B; scale bar in B”” is 200µm for A’ and B’ and 300µm for A”, B”, A’” and B’”.

### Reduction in GAD67-GFP positive cells in *Sema6A* KO mice

Given the distinct biochemical properties and intracellular distributions of GAD67 and GAD65 as well as the expression in distinct neuronal cell types [36–38], analysis of both GAD65+ and GAD67+ interneuron populations was required to fully evaluate the contribution of *Sema6A* toward GABAergic interneuron migration. Therefore GAD67-GFP knock-in mouse line [33] in which all GAD67+ inhibitory neurons express GFP were crossed with *Sema6A* mice. As previously reported in other studies [38], our results first showed that GAD67 appears to be the predominant GAD form in the neocortex (Fig. 3B’) compared to GAD65 expression (Fig. 2B’) in control mice at P30. It also appeared that the expression of GAD67, like GAD65, was affected by the loss of *Sema6A* gene. When compared to the control mean (referenced as 100% of the GAD67-GFP positive population), analysis of the robust GAD67-GFP within the *Sema6A* mutant mice revealed a reduction of 26% of GAD67-GFP+ cell number in the cortical area (Fig. 3A’ and 3C, mutant n=7, control n=12), 42% in the hippocampal area (Fig. 3A” and 3D, mutant n=7, control n=12), and 52% in the RTN area (Fig. 3A’” and 3E, mutant n=7, control n=12) at the adult stage (P30). Although all the brain areas that were analyzed displayed GAD67-GFP+ cell density loss, this loss was differentially affected depending on the brain area, with the RTN area affected the most. GABAergic cell organization was also affected in the RTN with the presence of holes in the mutant compared to the controls (Fig. 5C and C’, white arrows).

**Figure 3:**
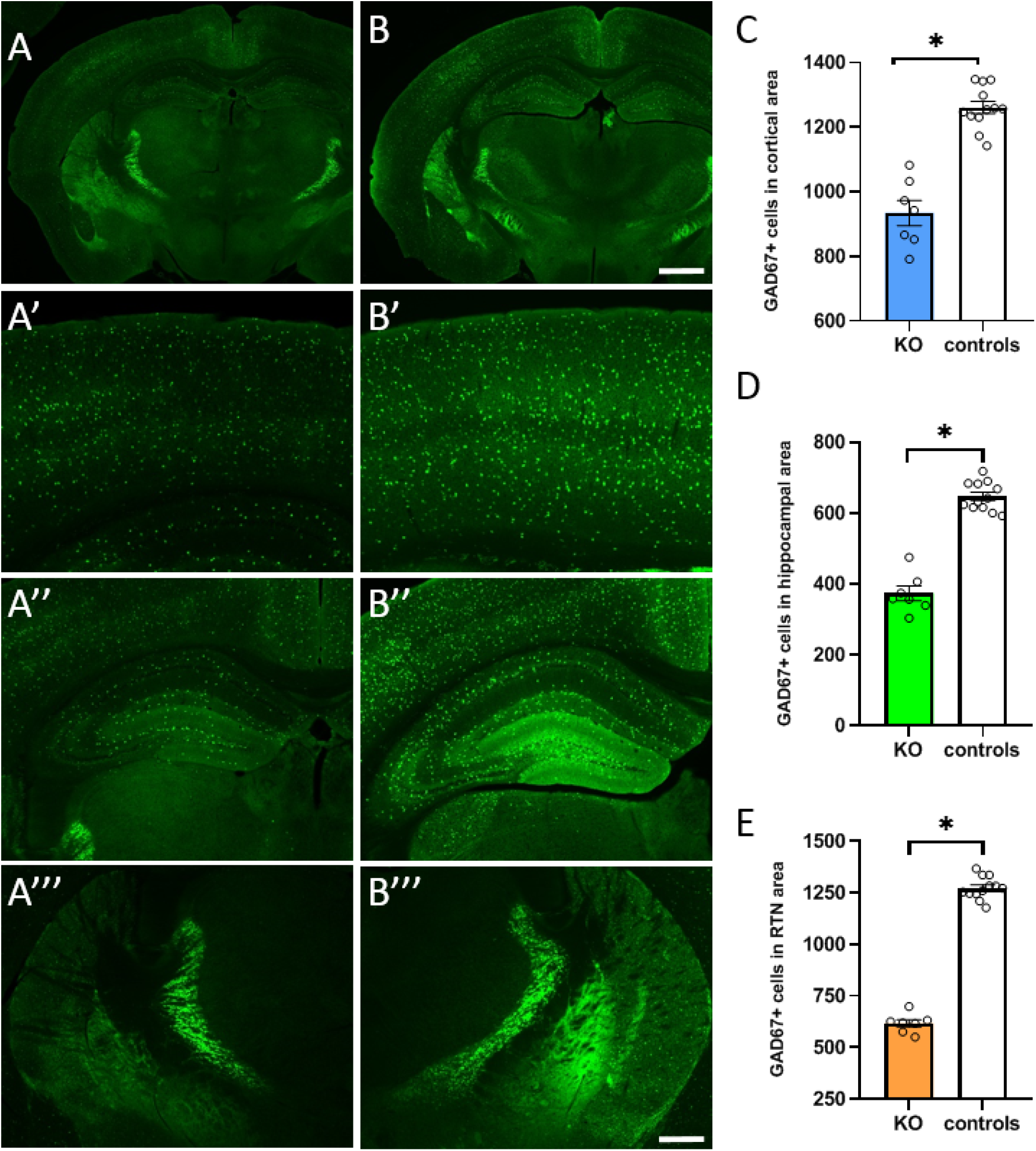
*Sema6A* mutant mice display a decrease in Gad67-GFP positive cells. GAD67-GFP mice were crossed with *Sema6A* mutant mice to allow for reliable tracking of the GAD67 interneurons population (green staining on panels A-A”’, B-B’”). Panels A’A’” shows GAD67-GFP positive cells (in green) in *Sema6A* mutant mouse at P30. A’, A” and A’” are higher magnification views of the cortex (S1), hippocampus, and RTN, respectively. Panels B, B’B’” shows GAD67-GFP positive cells in *Sema6A* control mouse at P30. B’, B” and B’” are higher magnification views of the cortex, hippocampus, and RTN areas, respectively. Panels C, D and E represent GAD67-GFP positive cell count with a reduction of GAD67 positive cells seen in *Sema6A* mutant mice at P30 in the cortical area (C, blue bar: KO, n=7, white bar: WT, n=12, p value<0.0001), hippocampal area (D, green bar: KO, n=7, white bar: WT, n=12, p value<0.0001), and reticular thalamic nucleus (E, orange bar: KO, n=7, white bar: WT, n=12, p value<0.0001). Scale bar in B is 1000µm for A and B; scale bar in B”” is 200µm for A’ and B’ and 300µm for A”, B”, A’” and B’”.

### PV Reduction in *Sema6A* KO mice

GABAergic interneurons have been shown to be highly heterogeneous and are classified into subpopulations using the expression of neurochemical markers. GABAergic interneurons may be classified by their expression of the calcium binding proteins Parvalbumin (PV), Calbindin (CaB), and Calretinin (CaR) (McDonald and Mascagni, 2001), with PV neurons being the most abundant (Cowan et al., 1990; Kawaguchi and Kubota, 1993). PV-expressing interneurons are characterized as fast spiking interneurons involved in gamma-oscillation generation that synchronize cortical activity during cognitive processing (Bartos et al., 2007; Sohal et al., 2009). To better identify the interneuron population affected by the loss of *Sema6A* gene, PV, CaB, and CaR protein expressions were analyzed. PV immunostainings at the adult stage (P30) revealed that PV expression is affected in the *Sema6A* mutant mice (Fig. 4C, mutant mice n=9) compared to controls (Fig. 4B, control mice n=10). Indeed a 46% cell density loss is seen in the cortical area (Fig. 4A’ and 4C) and although this cortical PV cell loss might appear more dramatic in the upper cortical layers, all cortical layers are affected by this PV interneuron loss. Cell count analysis also revealed a PV cell density loss of 45% in the hippocampal area (Fig. 4A” and 4D, mutant n=9, control n=10) and 37% in the RTN area (Fig. 4A’” and 4D, mutant n=10, control n=9). There were no differences between male and female *Sema6A* mutant or control mice (see methods for more details). This reduction in PV cell number was seen as early as PV expression turns on in the RTN area (Fig. 5A, A’, B and B’) at P7 when *Sema6A* mutant mice were compared to *Sema6A* control mice.

**Figure 4:**
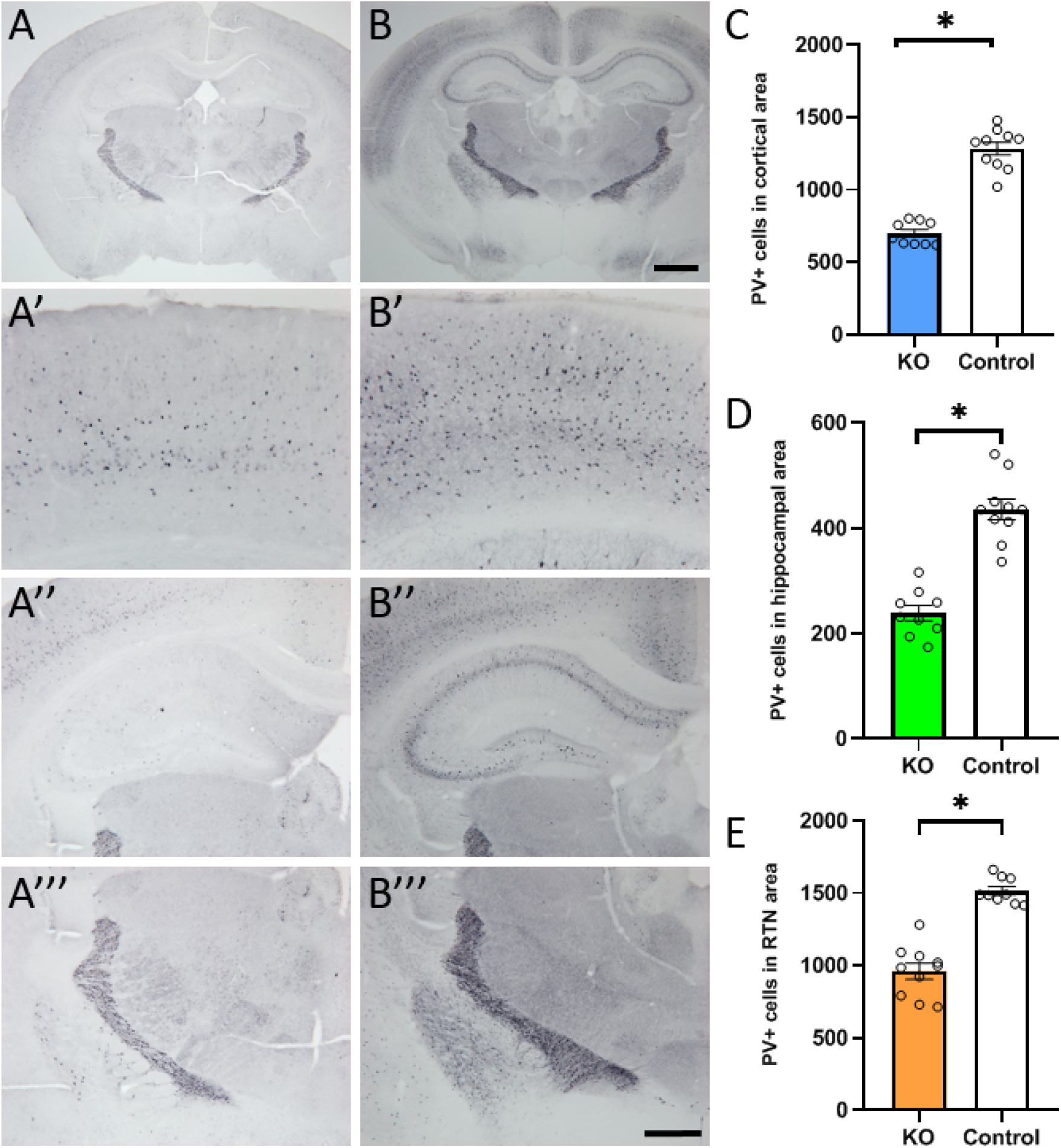
Decreased Parvalbumin cell number in *Sema6A* mutant mice. Parvalbumin staining was performed on P30 *Sema6A* mutant and control mice. Panels A, A’, A’” show PV cells in *Sema6A* mutant mouse at P30. A’, A” and A’” are higher magnification views of the cortex (S1), hippocampus, and RTN, respectively. Panels B, B’, B’” show PV cells in *Sema6A* WT mouse at P30. B’, B” and B’” are higher magnification views of the cortex, hippocampus, and RTN, respectively. Panels C, D and E display the reduction in PV positive cell count in *Sema6A* mutant mice at P30 in the cortical area (C, blue bar: KO, n=9, white bar: WT, n=10, p value<0.0001), hippocampal area (D, green bar: KO, n=9, white bar: WT, n=10, p value<0.0001), and reticular thalamic nucleus (E, orange bar: KO, n=9, white bar: WT, n=10, p value<0.0001). Scale bar in B is 1000µm for A and B; scale bar in B”’ is 200µm for A’ and B’ and 500µm for A”, B”, A’” and B’”.

**Figure 5:**
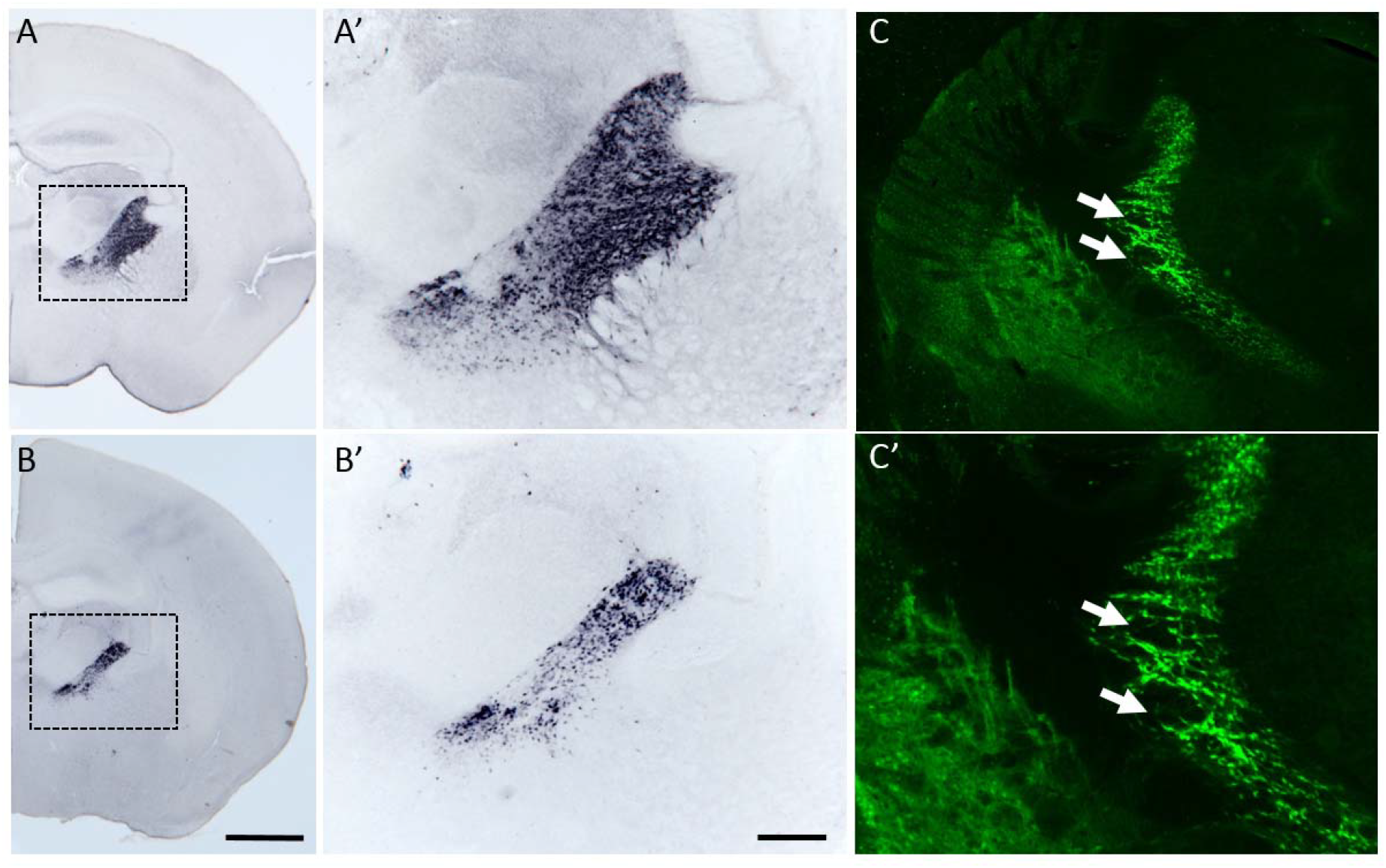
RTN GABAergic cell loss in *Sema6A* mutant mice. Parvalbumin staining was performed on P7 *Sema6A* mutant and control mice and was found in the RTN area. Panels A and A’ show PV cells in *Sema6A* control mouse. A’ is a higher magnification view of the RTN area. Panels B and B’ show PV cells in *Sema6A* mutant mouse. B’ is a higher magnification view of the RTN area. At this stage the PV cell reduction can be seen in the *Sema6A* mutant (B) compared to the Sema6 control mice (A). Panels C and C’ display an example of the GABAergic cell reduction in the RTN, where holes (white arrows in both C and C’) can be seen in some areas of the RTN in *Sema6A*/GAD-67 GFP mutant mice at P30. Scale bar in B is 1000µm for A and B; scale bar in B’ is 300µm for A’, B’ and C’ and 150µm for C’.

### Both Calretinin and Calbindin expressions are not affected in *Sema6A* KO mice

It is well recognized that PV, CaR, and CaB are useful markers for categorizing interneuron populations as they are expressed by more than 80% of the GABAergic neurons [39–44]. As this study aimed to investigate the contribution of *Sema6A* toward GABAergic interneuron populations, along with PV, CaR and CaB expressing interneurons were analyzed in *Sema6A* mutant mice (Fig. 6 and Fig. 7, respectively). Results showed that the expression of CaR was unaffected by the loss of the *Sema6A* gene (Fig. 6). When both primary somatosensory cortical and hippocampal areas were analyzed at P30, there were no significant differences in CaR expression in the brain areas when *Sema6A* mutant mice (Fig. 6 A, A’, A’”) were compared to *Sema6A* control mice (Fig. 6B, B’, B’”). Expression of CaR in the RTN brain area was not analyzed since RTN is comprised of 98% of PV interneurons [45]. Cell count analysis was performed in both the cortical (S1) and hippocampal areas and revealed no significant differences (data not shown). Similarly to the PV expression, CaB expression was analyzed in the same brain areas, except for the RTN area. When cell count analysis was performed there were no differences in both the primary somatosensory cortical (Fig. 7 A, A’ *Sema6A* mutant mice compared to Fig. 7B, B’ control mice) and hippocampal (Fig. 7 A, A” *Sema6A* mutant mice compared to Fig. 7B, B” control mice) interneuron cell populations expressing CaB. Taken together these results suggest that of the GABAergic calcium-binding interneurons the loss of *Sema6A* affects only the PV interneuron population.

**Figure 6:**
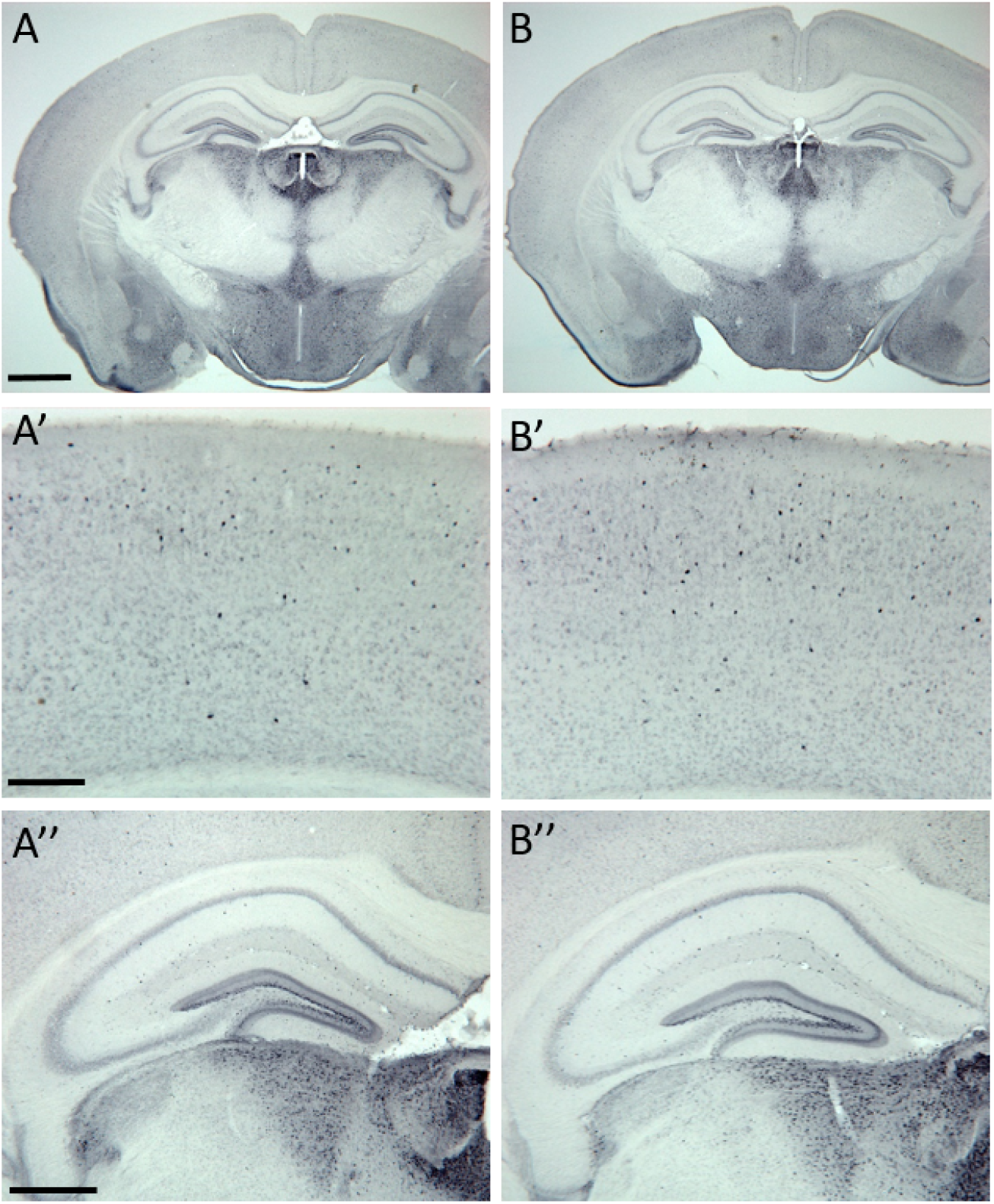
*Sema6A* mutant mice display no alteration in Calretinin interneuron population. Calretinin staining was performed on coronal brain sections at P30. No differences in expression were observed between the *Sema6A* mutant mice (A, A’ is a high power view of the cortical area, A” is a high magnification of the hippocampal area) and the *Sema6A* control mice (B, B’ is a high power view of the cortical area, B” is a magnification of the hippocampal area). Cell density analysis in the cortical (S1), hippocampal, and RTN areas revealed no significant difference in CaR interneurons (data not shown) between *Sema6A* mutant and control mice. Scale bar in A is 1000µm for A and B; scale bar in A’ is 200µm for A’ and B’, scale bar in A” is 500µm for A”, B”.

**Figure 7:**
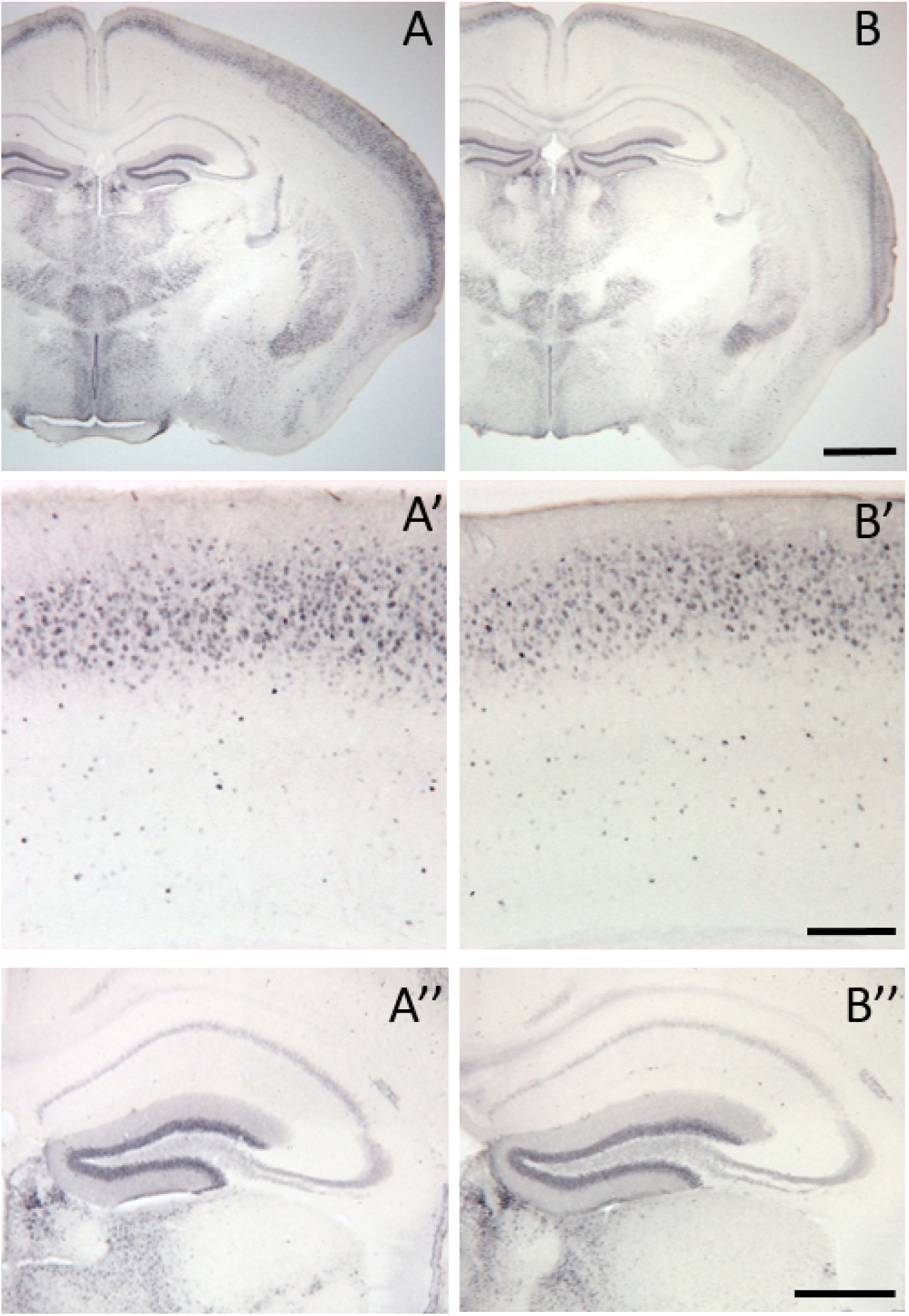
Calbindin interneuron population was not affected in *Sema6A* mutant. Calbindin staining was performed on coronal brain sections at P30. No differences in expression were found between the *Sema6A* mutant mice (A, A’ is a high power view of the cortical area, A” is a high magnification of the hippocampal area) and the *Sema6A* control mice (B, B’ is a high power view of the cortical area, B” is a magnification of the hippocampal area). Cell count analysis in the cortical (S1), hippocampal, and RTN areas revealed no significant difference in the CaB interneuron population (data not shown) between *Sema6A* mutant and control mice. Scale bar in B is 1000µm for A and B; scale bar in B’ is 200µm for A’ and B’, scale bar in B” is 500µm for A”, B”.

## DISCUSSION

In the present study we took advantage of genetically modified mouse lines to examine the GABAergic interneuron population when *Sema6A* gene is knocked out. Knock-in mouse lines expressing either GAD65-GFP or GAD67-GFP were crossed with *Sema6A* mouse line to examine GABAergic interneuron populations. The advantage of using these two GFP reporter lines is a strong, robust and reliable staining of either GAD65 or GAD67 which facilitates our analysis of the GABAergic interneuron populations in the *Sema6A* mutant mouse line compared to GABA or GAD65 or GAD67 immunostainings. Our major findings using these genetic manipulations are as follow: 1. Loss of *Sema6A* gene leads to a dramatic overall GABAergic cell reduction in the primary somatosensory cortical, hippocampal, and RTN brain areas. To our knowledge this is the first study reporting a GABAergic interneuron loss in the RTN, an area known to play a role in the brain’s attentional network. 2. Analysis of the subpopulations of inhibitory neurons revealed that the fast spiking interneurons, PV, are the only GABAergic interneurons affected by the *Sema6A* mutation as CaR and CaB cell populations remain unchanged. 3. Loss of the PV interneuron cell population can be observed in the RTN as early as the first postnatal week, when PV expression is initiated.

### *Sema6A* loss leads to GABAergic cell loss

In this study, we aimed to understand E/I imbalance at a cellular level, and hypothesized that either the production, migration, cell-type specification, or maturation of GABAergic interneurons may be affected in ASD. To this end, *Sema6A* gene contribution was analyzed to understand whether the loss of this axon guidance molecule can affect GABAergic interneuron populations. Our results showed a reduction in GABAergic cell number in the primary somatosensory cortical, hippocampal, and RTN areas, thereby revealing the importance of *Sema6A* in the formation of GABAergic neuronal networks in these brain areas. Other studies have previously documented the role of some semaphorin family members in interneuron cell migration during brain development. Sema3A knockout mice display a reduced number of interneurons in the developing cortex due to altered migration [46–49]. These studies focused only on embryonic stages and did not indicate adult GABAergic interneuron fate, nor which interneuron cell type was affected as a result of the migration defect. *Sema3A* is not the only member of the semaphorin family to control cortical interneuron migration [46–49] and cerebellar interneuron branching [50] as studies have described the role of *Sema6A* in cerebellar granule cell migration [30, 51]. *Sema3C*, another member of the family, also plays a role in GABAergic fate, as it is implicated in the transient presence of GABAergic interneurons during the formation of the corpus callosum (CC) [52]. It is interesting to note that the loss of these transient GABAergic interneurons in the CC resulted in major pathfinding defects in the CC [52]. Such defects in CC formation were also reported in *Sema3A* knockout mice [33, 53] and provide more evidence for a role of semaphorin members in ASD as 1/3 of individuals with autism have an abnormal CC [54]. While most studies have focused on the importance of semaphorins in the control of neuronal migration and axon guidance during brain development, recent studies have reported the role of semaphorins in the maturation of cortical circuits [55, 56]. Indeed failure in GABAergic circuitry formation was described in the *Sema7A* knockout mice, however the GABAergic cell loss was only reported in layer 4 of the barrel cortex [55] compared to our results showing that all the cortical layers are affected in the *Sema6A* mutation. More recently, Greg Barnes’ group published an informative study on interneuron-specific knockout mouse of *Sema3F* [56]. Their data showed a 20% reduction of GABAergic cells in the somatosensory cortex while we report a 50% reduction of GAD65-GFP positive cells and 26% reduction of GAD67-GFP positive cells in the same cortical area. The differences in cell number reduction between the 2 studies might be explained by the fact that *Sema6A* and *Sema3F* play different roles in either the GABAergic cell production, migration or maturation. *Sema7A* knockout mice also displayed a GABAergic cell loss in the hippocampus, however it was specifically located in the CA1 area [56] whereas our data showed that in *Sema6A* mutant mice all the hippocampal sub-regions had a reduction in GABAergic cell number. Taken together, our data suggest the importance of *Sema6A* gene in either the production, proliferation and/or migration of GABAergic interneurons since the disruption of this gene leads to a GABAergic cell loss in several brain areas.

### Loss of *Sema6A* gene leads to a loss of PV cells

Of the GABAergic interneurons, PV neurons are the most abundant [57, 58] and are involved in the generation of cortical gamma-oscillations that synchronize cortical activity driving cognitive processing [59, 60]. Abnormalities of PV neurons have attracted attention as one of the potential causes underlying neuropsychiatric disorders [11]. Studies on postmortem brains of individuals with schizophrenia, autism, and bipolar disorder have revealed that the number of PV neurons is lower in the frontal cortex, entorhinal cortex, and hippocampus [44, 61–66]. Several studies using genetic mouse models to investigate the etiology of ASD also reported PV defects in these animal models [21, 67–78]. As we seek to understand the mechanisms behind human GABAergic defects in ASD, *Sema6A* knockout mice were crossed with either a knock-in mouse model expressing GAD65-GFP or GAD67-GFP. Since our results showed a loss of GABAergic interneurons in *Sema6A* knockout mice, we analyzed the expression of different populations of the GABAergic interneurons and found that the PV cell population displayed a reduction in cell number, whereas CaR and CaB cell populations were unaffected by the mutation. Similar to our observations, in the FMR1 knockout mice. GABAergic cell loss was only targeted to the PV cell population, as no changes in either CaR or CaB were seen [72]. FMR1 knockout mice displayed a 20% reduction in PV cell populations in the somatosensory cortex [72], while our results showed a more pronounced PV cell density loss with a 46% reduction in the primary somatosensory cortical area. Hippocampal PV cell reduction was also reported in the Neuropilin-2 mutant mouse, with a 42% PV cell reduction [21] compared to the 45% PV cell reduction seen in the *Sema6A* mutant mice. Since Neuropilin 2 has been shown to interact with *Sema6A* [79, 80], these very similar PV cell losses seen in both *Sema6A* and Neuropilin 2 knockout mice could suggest that *Sema6A* interacts with Neuropilin 2 to facilitate the migration of the GABAergic interneurons, and disruption of either leads to alterations of the GABAergic interneurons and more specifically the PV population. CNTNAP2 knockout mice, another well-known ASD model, also displayed a 20% PV cell loss in both the cortical and hippocampal areas [71]. This PV cell reduction was lower than the one we observed in the *Sema6A* mutant mice, which displayed a 46% and a 45% reduction in the cortical and hippocampal areas, respectively, suggesting that there are different mechanisms controlling the GABAergic interneurons in these two ASD animal models. In the Neuroligin-3 mutant mice, the cortical PV cell loss [68] was very similar to the one we observed in the *Sema6A* knockout mice (~50%), however no PV cell loss was reported in the hippocampal area, indicating that Neuroligin-3 might only play a role in the fate of cortical GABAergic interneurons, whereas *Sema6A* might be involved in both hippocampal and cortical GABAergic cell fate. Taken together, it appears that similar to other mouse models that previously reported a PV cell reduction, *Sema6A* mutant mice also display a PV cell loss, suggesting that in ASD mouse models PV defects could be a common feature. Recent studies on Shank1 and also Shank3 mutant mice suggested the PV reduction observed in both knockout mouse models was due to a PV downregulation rather than a neuronal loss [67, 69, 81, 82]. Based on our results showing an overall GABAergic cell loss, using GAD transgenic mouse lines to reliably visualize GABAergic interneurons, and the fact that neither CaR nor CaB interneurons are affected, we believe that the PV loss that we see in the *Sema6A* mutant mice is due to a PV cell loss rather than a downregulation of PV.

### RTN brain area displayed the greatest GABAergic cell loss in *Sema6A* mutant mice

The RTN, discovered by Kölliker [83] and located between the internal capsule and the external medullary lamina, originates from the ventral thamalus [84] and receives inputs from the cerebral cortex and dorsal thalamic nuclei [85, 86]. It is sub-divided into several sectors [87–89] (1 limbic, 1 motor and 5 sensory sectors, including visual, auditory, somatosensory, gustatory and visceral) and GABAergic neurons appear to be the major RTN cell type [90, 91], with 98% of these GABAergic neurons expressing PV [45]. RTN has been described as a gate keeper [92] and has been shown to play a role in attention [93–97], multisensory gating [98], emotions [97], sleep [99, 100], and pain regulation [101]. More recent work by Halassa’s group [102] described the RTN as a switchboard regulating the information received by the brain, and more importantly filtering out unnecessary information, thereby reinforcing the importance of the RTN in attention processes and the consequences that might arise if such neural circuits become defective. While Halassa’s group also investigated the role of *Ptchd1*, a gene mutated in ASD (~1% of all individuals with ASD), and reported RTN impairments using conditional *Ptchd1* knockout mouse with attention deficit and hyperactivity, they did not report any loss of any GABAergic interneurons in this brain area [103]. The only PV cell loss that has been reported in the RTN area was found in subjects with either schizophrenia or bipolar disorder [104]. Although GABAergic cell loss and more specifically PV cell loss was reported in several studies using ASD mouse models, the interneuron loss was either seen in the cortical area and/or the hippocampus. Therefore to our knowledge this is the first study reporting a GABAergic interneuron cell loss in the RTN when an axon guidance gene reported as an ASD candidate gene is mutated. The reduction of PV cell number (37%) observed in the RTN of *Sema6A* mutant mice suggests that *Sema6A* might play a role in the RTN formation by controlling the migration of the GABAergic interneurons into that area. *Sema6A* may have a deep impact in the RTN function since such a cell loss might contribute to cognitive, emotional, attentional, sensory, and sleep defects, all of which have been observed in individuals with ASD.

## LIMITATIONS

*Sema6A* mutant mice have been a valuable tool to assess the potential mechanisms altered in ASD, however it is important to acknowledge the limitation of animal models and the degree to which they can match the complexity of the human brain.

## CONCLUSIONS

### *Sema6A* gene and its implication in E/I balance

The goal of our study was to focus on the potential role of *Sema6A* in ASD and assess whether the loss of an axon guidance molecule could affect the proper formation of neuronal circuits by disrupting the GABAergic interneuron population. We did not intend to characterize the *Sema6A* mutant mice phenotype, since axon guidance, cell migration, and behavioral defects have been previously reported by other groups [27–30, 51, 105–109], but rather we intended to focus on whether *Sema6A* mutants can be used as an informative model to investigate the E/I imbalance. As our results showed GABAergic cell loss, and more specifically PV cell loss, in key brain areas involved in ASD, our study revealed the implication of *Sema6A* E/I imbalance and its role on neurodevelopmental defects/ASD. Indeed, *Sema6A* mutant mice also displayed cortical lamination defects in the auditory cortex (data not shown, [24]) similar to the disorganization of the cortical layers reported in individuals with autism [7], adding evidence to the role of *Sema6A* in ASD. How PV cells are being disrupted in ASD still remains unclear, however it would appear that since only PV interneurons are affected by the loss of *Sema6A*, *Sema6A* is very likely to play a role in the medial ganglionic eminence (MGE) maturation rather than the lateral (LGE) or caudal ganglionic eminence (CGE) maturation which are producing CaR interneurons. Further studies looking at brain development and the generation of interneurons in *Sema6A* mutant mice will help to elucidate whether the proliferation, migration, or specification of PV interneurons is affected as a mechanism contributing to human GABAergic disruptions in ASD.

## LIST OF ABBREVIATIONS

ASD: autism spectrum disorder
caB: Calbindin
caR: Calretinin
CGE: caudal ganglionic eminence
E/I: excitation/inhibition
GAD: Glutamate decarboxylase
GFP: Green fibrillary protein
LGE: lateral ganglionic eminence
MGE: medial ganglionic eminence
MRI: magnetic resonance imaging
PV: Parvalbuim
RTN: reticular thalamic nucleus
Sema6A: semaphoring 6A

## DECLARATIONS

## Acknowledgements

We would like to thank Dr. John Hussman, Dr. Gene Blatt, and Ms. Elizabeth Benevides for critical reading and editing of the manuscript, and Serena Edwards for technical assistance. We would like to acknowledge Dr. Kevin Mitchell, Dr. Giovanna Tosato, and Dr. Ombretta Salvucci for the *Sema6A* mutant mice, as well as Dr. Alain Chedotal, Dr. Alex Kolodkin, and Dr. Adam Puche for helpful discussions. We thank Dr. Louis DeTolla from the University of Maryland Baltimore for veterinary and consulting services.

## Funding

This project is supported by the Hussman Foundation Grant #HIAS15005 (CP).

## Availability of data and materials

Datasets for the current study can be made available from the corresponding author on reasonable request.

## Authors’ contributions

CP conceived the study and secured the funds. KM and CP performed the animal studies, analyzed statistics and wrote the manuscript. GS, YY and TC assisted in the experiment design. All authors have read and approved the final manuscript.

## Ethics approval

Animals: the use of animals was in accordance with the Institutional Animal Care and Use Committees (IACUC) from the Hussman Institute for Autism and University of Maryland, Baltimore (UMB).

## Consent for publication

Not applicable

## Competing Interests

The authors declare no competing interests.

